# Chromosomal instability mediates immune exclusion and response to cytotoxic chemotherapy in colorectal liver metastases

**DOI:** 10.1101/2021.09.22.459429

**Authors:** Carlos A Martinez, Liam F Spurr, Soumya C Iyer, Sian A Pugh, John A Bridgewater, John N Primrose, Enric Domingo, Timothy S Maughan, Michael I D’Angelica, Mark Talamonti, Mitchell C Posner, Philip P Connell, Ralph R Weichselbaum, Sean P Pitroda

## Abstract

The genomic drivers of immune exclusion in colorectal cancer liver metastases (CRCLM) remain poorly understood. Chromosomal instability (CIN), resulting in aneuploidy and genomic rearrangements, is the central pathway of mismatch repair-proficient colorectal cancer pathogenesis; however, it is unknown whether CIN impacts the outcomes of patients with limited spread of CRCLM treated with curative intent cytotoxic chemotherapy and surgery. Herein, we examined the relationship between CIN and the molecular subtypes of CRCLM, immune signaling, treatment sensitivity, and patient outcomes in three independent CRCLM patient cohorts. We established that a previously developed 70-gene CIN signature (CIN70) is a reliable measure of CIN, encompassing features of both aneuploidy and cellular proliferation. We demonstrated that tumors with the canonical subtype of CRCLM exhibit elevated levels of CIN and aneuploidy. Genomically unstable tumors were associated with an immune-depleted tumor microenvironment, and patients with genomically unstable tumors were at increased risk for disease progression in adverse metastatic sites, resulting in poor progression-free and overall survival. However, high-CIN tumors were particularly susceptible to DNA-damaging chemotherapies, including topoisomerase inhibitors, as well as radiation therapy. Treatment with genotoxic agents depleted CIN-rich cell populations, which resulted in a concomitant increase in intratumoral CD8+ T-cells in patients with primary rectal, breast, and bladder cancer. Taken together, we propose a mechanistic explanation for why cytotoxic chemotherapy can augment anti-tumor immunity and improve outcomes in patients with genomically unstable cancers.

## Introduction

A subset of patients with metastatic colorectal cancer experiences long-term survival following curative-intent treatment with chemotherapy and surgical removal of primary tumors and limited metastases. However, it remains difficult to predict which patients will derive benefit from these potentially morbid interventions. We previously conducted a multidimensional computational analysis to define three robust integrated molecular subtypes of mismatch repair-proficient (microsatellite stable, MSS) colorectal cancer liver metastases (CRCLM), which we designated as canonical, immune, and stromal subtypes*(1)*. Patients with the immune subtype of CRCLM and favorable clinicopathologic features experienced extended survival rates, and many patients were cured. By contrast, the canonical subtype of CRCLM exhibited decreased expression of immune and stromal signatures in the context of activated E2F/MYC and aberrant cell cycle checkpoint pathways. However, the molecular mechanisms driving differential outcomes between these subtypes remain poorly understood.

Chromosomal instability (CIN) is the central driver of genomic instability and tumor development in sporadic, MSS colorectal cancer*(2)*. CIN, defined as a high rate of whole or segmental chromosomal copy number aberrations, can be subdivided into numerical CIN, which involves differences in chromosome number, and structural CIN, which involves chromosomal rearrangements, deletions, and insertions*(3)*. Importantly, CIN leads to elevated intratumoral tumor heterogeneity (ITH) and aneuploidy, a state of aberrant chromosome or chromosome arm number, which are associated with altered responsiveness to treatment and increased probability of metastases*(4)*. Nevertheless, it remains unclear whether CIN is directly associated with clinical outcomes for patients treated for CRCLM.

In this study, we performed a multi-omic investigation to examine the functional consequences of CIN in CRCLM. We leveraged three large clinical cohorts of extensively annotated patients (total n=336) with MSS CRCLM without extrahepatic metastases treated with cytotoxic chemotherapy and complete resection of primary and metastatic tumors from two retrospective cohorts and one prospective clinical trial (**Table 1**)*(1, 5, 6)*. Herein, we report that the immune-depleted canonical subtype of CRCLM is enriched for CIN and that the degree of CIN is independently prognostic for adverse clinical outcomes. Furthermore, we extended these findings by demonstrating in primary rectal, breast, and bladder cancers that cytotoxic therapies deplete CIN-rich cell populations within tumors, thereby augmenting intratumoral adaptive immunity and improving patient outcomes.

**Table 1.** Colorectal Cancer Liver Metastasis Patient Cohorts.

| **Clinical Cohorts** | **Type** | **Size** | **Analyses** | **Clinical Endpoints** |
| --- | --- | --- | --- | --- |
| UCMC<sup>1</sup> | Retrospective cohort | 93 | Gene expression, mutations & CNV | DFS, OS & POF |
| MSKCC<sup>5</sup> | Retrospective cohort | 96 | Gene expression | DFS, OS & POF |
| UK/New-EPOC<sup>6</sup> | Randomized control trial | 147 | Gene expression, mutations & CNV | PFS & OS |

CNV: copy number variation; DFS: disease free survival; PFS: progression free survival; OS: overall survival; POF: patterns of failure

## Results

### CIN is prognostic for survival in colorectal liver metastasis

Given the importance of CIN as a driver of cancer progression in mismatch repair-proficient colorectal cancer, we investigated the impact of CIN on clinical outcomes after curative-intent treatment of CRCLM with cytotoxic chemotherapy and surgical resection of primary tumors and limited metastases. Patients in the UCMC and MSKCC were well-matched in terms of clinical and pathological factors; however, patients in the MSKCC cohort were more likely to undergo surgical resection of a greater number of liver metastases. By contrast, patients in the UK/New-EPOC cohort were slightly older and more likely to present with a shorter disease-free interval between primary tumor and metastatic diagnoses, as well as a larger number of liver metastases (**Supplementary Table S1**). We quantified CIN by utilizing a gene expression-based signature of 70 genes (termed CIN70) which strongly correlates with functional aneuploidy in tumors and is prognostic of clinical outcomes in 12 cancer data sets representing 6 different cancer types*(7)*. Interestingly, metastatic tumors harbor higher CIN70 scores than do primary tumors, suggesting that CIN may help promote the formation of metastases*(7)*.

We found that canonical CRCLMs exhibit higher CIN70 scores than do immune and stromal CRCLMs (means = 0.45 for canonical, 0.31 for immune, and 0.26 for stromal subtypes, respectively; one-way ANOVA p < 0.0001) (**Figure 1A)**. CIN70 scores were independent of established adverse clinical and pathological prognostic factors that define the Clinical Risk Score (CRS) in CRCLM*(8)*, including the presence of more than one liver metastasis, lymph node positivity, carcinoembryonic antigen (CEA) greater than 200 ng/mL, disease-free interval between primary tumor diagnosis and development of liver metastasis less than one year, and liver metastasis size greater than 5.0 cm (**Supplementary Figure S1A**).

**Figure 1.**
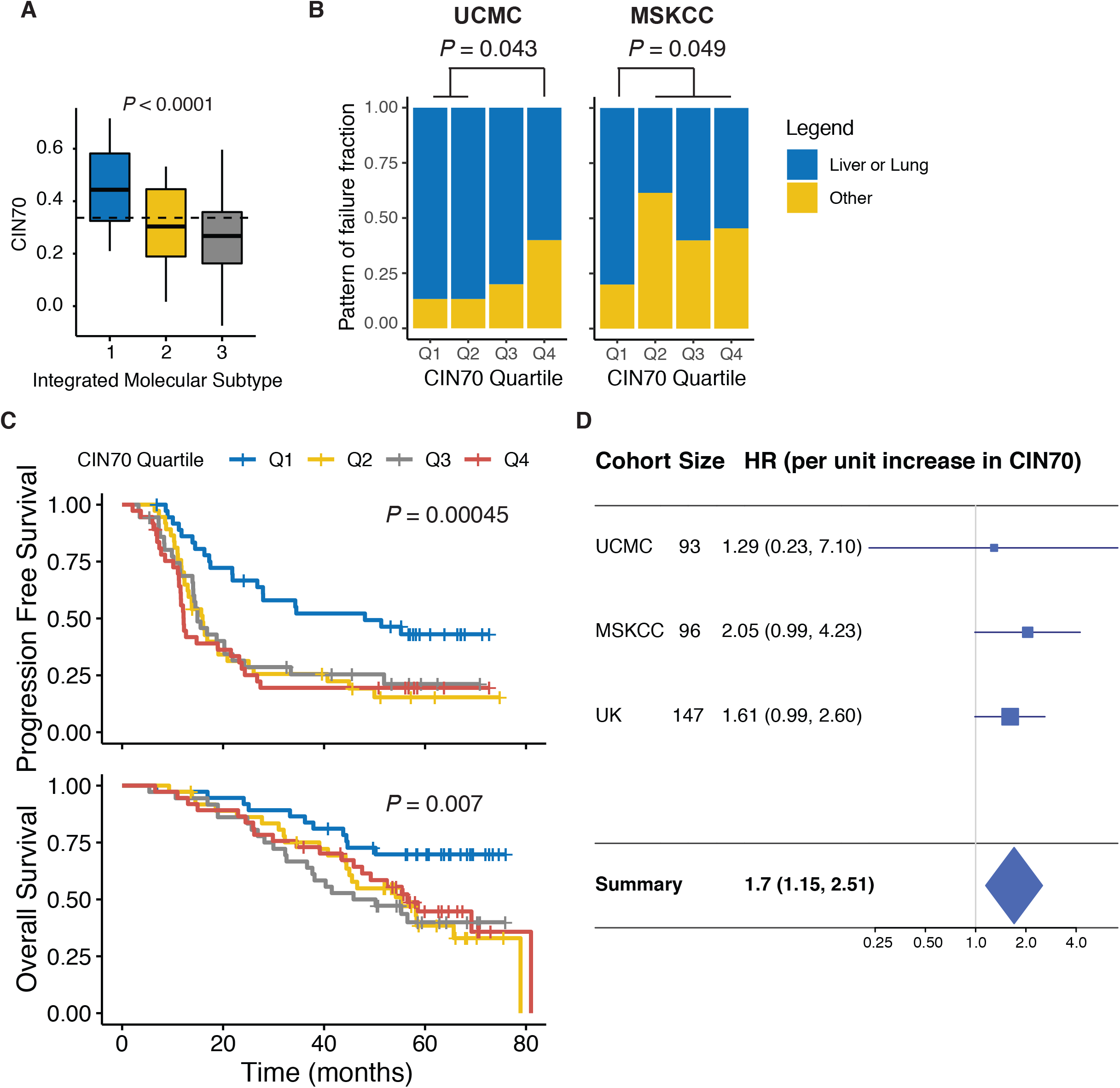
CIN is prognostic for survival in colorectal liver metastasis. **A)** Boxplots of CIN70 scores as a function of the three integrated molecular subtypes in the UCMC dataset where subtypes 1, 2, and 3 correspond to canonical, immune, and stromal subtypes, respectively; dashed line denotes the average CIN70 score of all samples; p-value was calculated using a one-way ANOVA test (n = 93). **B)** Stacked barplots showing the percentage of patients with liver or lung metastatic recurrence (favorable; blue) and the percentage of patients with metastatic recurrences to other sites (unfavorable; yellow) following initial curative-intent treatment of primary and metastatic CRC liver metastases; the vertical axis shows the fraction of patients corresponding to each pattern of failure; the horizontal axis denotes the CIN70 quartile; the left panel corresponds to the UCMC dataset (n = 60), right panel corresponds to the MSKCC dataset (n = 49); p-values were calculated using a chi-square test. **C)** Kaplan-Meier curves of the UK/New-EPOC dataset for progression free survival (upper panel) and overall survival (lower panel); line colors correspond to CIN70 quartiles; p-value for statistical difference was calculated using the log-rank method (n = 147). **D)** Meta-analysis of the univariate hazard ratios of overall survival for CIN70 scores; horizontal blue lines denote the 95% CI; standard errors were calculating assuming random effects; horizontal axis shows the hazard ratio on a log scale; the boxes show the effect estimates from the single studies, while the diamond shows the pooled result (n = 93, 96, and 147 for UCMC, MSKCC, and UK/New-EPOC respectively).

Some studies suggests that high CIN is associated with an increased probability of metastases*(9)*. In our cohort of definitively treated metastases, patients with high CIN70 metastases were more likely to experience subsequent metastatic recurrences in adverse organ sites (i.e. beyond the liver and lung), including the brain, bone, and distant lymph nodes, as compared to low CIN70 metastases (OR = 4.3, Chi squared p = 0.043 for UCMC; OR = 4.0, Chi squared p = 0.049 for MSKCC) (**Figure 1B**). Importantly, metastatic recurrence beyond the liver and lung was consistently associated with a higher risk of death following treatment (HR = 2.40, univariate Cox p = 0.016 for UCMC; HR = 1.96, univariate Cox p = 0.029 for MSKCC) (**Supplementary Figure S1B**). In concert with these results, our analysis of the randomized UK/New-EPOC trial demonstrated that the lowest quartile of CIN70 was associated with favorable progression-free (HR = 0.43 [0.26 – 0.70, 95% CI], univariate Cox p = 0.00045) and overall survival (HR = 0.42 [0.22 – 0.81, 95% CI], univariate Cox p = 0.007) following metastasis-directed treatment when compared to the upper three quartiles of CIN70 (**Figure 1C**). Furthermore, a meta-analysis using univariate Cox proportional hazards analysis of the three cohorts showed that a per-unit increase in CIN70 was associated with a 1.7-fold ([1.15 – 2.51, 95% CI], univariate Cox p = 0.0074) risk for death (**Figure 1D)**. CIN70 remained independently prognostic of overall survival (HR = 1.57 [1.07 – 2.31, 95% CI], p = 0.02) in a multivariable Cox regression meta-analysis combining CRS and CIN70 (**Supplementary Figure S1C**). Taken together, these findings demonstrated that elevated CIN70 is associated with an increased propensity for metastatic dissemination and unfavorable survival in patients with CRCLM.

### CIN is associated with tumor aneuploidy and increased proliferation

Since CIN70 is a surrogate for functional aneuploidy*(7)*, we next investigated the relationship between CIN70 and tumor aneuploidy in CRCLM. Genomic copy number alteration (CNA) profiles were determined for a subset of CRCLMs in the UCMC cohort *(1)* that had been genomically characterized (OncoPlus assay; n = 59) and validated using an independent Memorial-Sloan Kettering Cancer Center (MSKCC) cohort (MSK-IMPACT assay; n = 315 non-MSI patients). The ASCETS algorithm was used to call arm-level somatic copy number alterations (aSCNAs) in each cohort*(10)*. The UCMC and MSKCC datasets exhibited a similar profile of amplifications and deletions across all chromosomes (**Supplementary Figure S2A**). In addition, a principal component analysis (PCA) demonstrated that the aSCNA profiles of the datasets did not reveal major differences between the two cohorts (**Supplementary Figure S2B**).

We then computed aneuploidy scores from the aSCNA calls as the fraction of called arms in a sample harboring an amplification or deletion. As expected, CRCLMs exhibited a marked increase in aneuploidy score when compared to matched adjacent histologically non-tumor liver specimens (**Supplementary Figure S2C**). Furthermore, the lowest quartile of CIN70 scores was observed in the tumors with the lowest aneuploidy in CRCLMs (mean Q1 = 0.24 vs. mean Q2-4 = 0.40, two-tailed t-test p = 0.025) (**Figure 2A**). In the UK CRCLM cohort, CIN70 also correlated with the fraction of genome altered (FGA; a measure of global somatic CNA) (mean Q1 = 6.30 vs. mean Q2-4 = 6.60, two-tailed t-test p = 0.0013) (**Figure 2B**) and number of coding mutations (mean Q1 = 6.30 vs. mean Q2-4 = 6.60, two-tailed t-test p = 0.0022) (**Figure 2C**).

**Figure 2.**
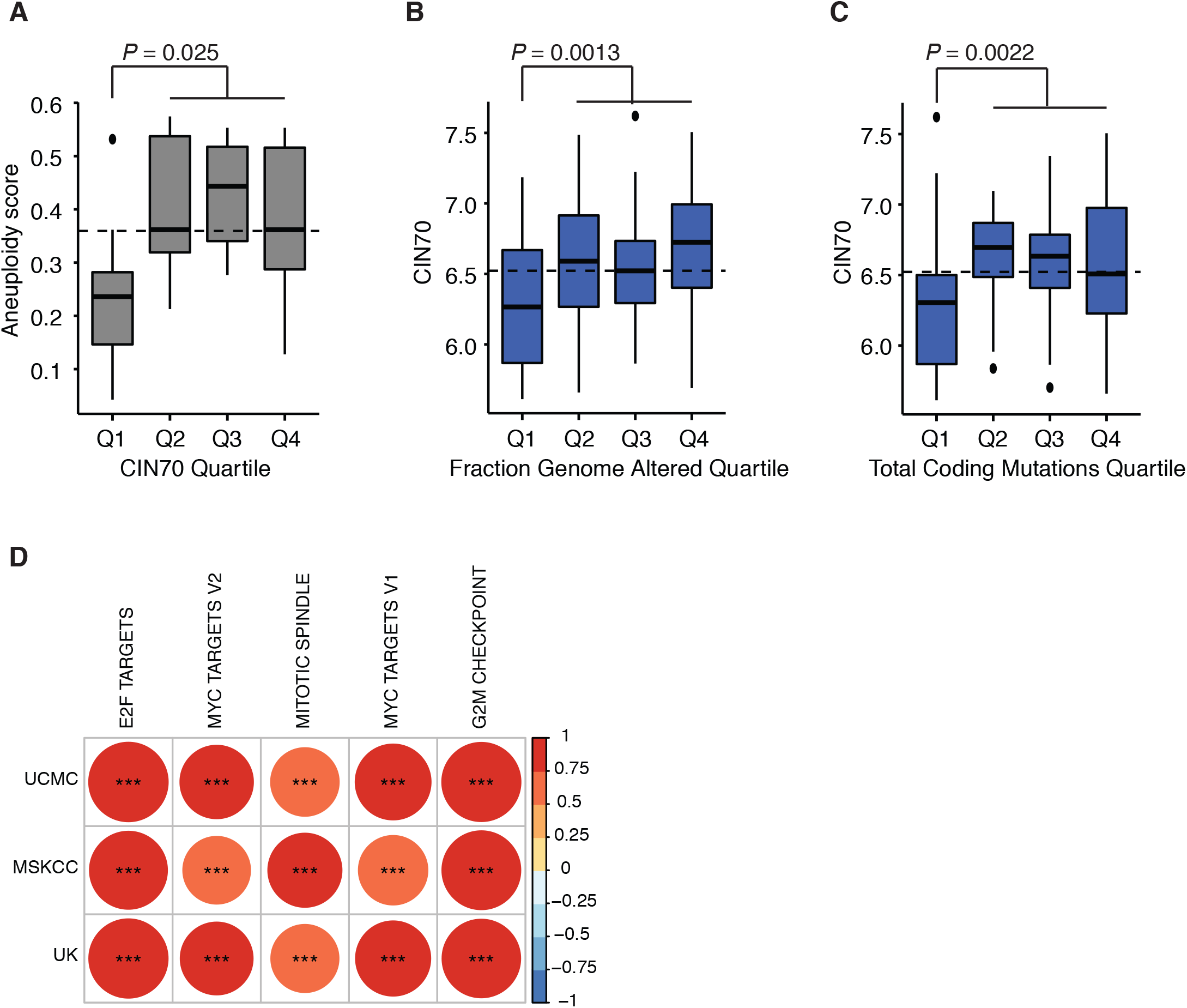
CIN is associated with tumor aneuploidy and increased proliferation. **A)** Boxplot of aneuploidy scores by CIN70 quartiles in UCMC dataset; dashed line corresponds to the mean aneuploidy score across all samples; p-value was calculated using a two-tailed t-test (n = 32). **B)** Boxplot of CIN70 scores by the fraction genome altered quartile in UK/New-EPOC dataset; p-value was calculated using a t-test (n = 147). **C)** Boxplot of CIN70 scores by the total coding mutations quartile in UK/New-EPOC dataset; p-value was calculated using a two-tailed t-test (n = 147). **D)** Correlation plot showing the Pearson correlations between ssGSEA scores for the Hallmark proliferation pathways and CIN70 scores among the three cohorts (UCMC, MSKCC, and UK/New-EPOC); circle size and color are proportional to the Pearson correlation; red denotes a positive correlation, while blue reflects a negative correlation; *P < 0.05, **P < 0.01, and ***P < 0.001 (n = 93, 96, and 147 for UCMC, MSKCC, and UK/New-EPOC respectively).

Recent studies have suggested that CIN is associated with increased tumor cell proliferation, which has been hypothesized to be mediated by the selective amplification of core regulators of proliferation*(11)*. We examined the correlation between CIN70 and the expression of cell cycle and proliferation pathways by correlating CIN70 scores with the results of Gene Set Enrichment Analysis (ssGSEA) for 50 Hallmark gene sets from the MSigDB. This demonstrated strong positive correlations (Pearson correlation p < 0.0001) between CIN70 and cell proliferation pathways across the three cohorts (**Figure 2D**). Additionally, high aneuploidy scores were correlated with increased expression of cellular proliferation pathways such as the Hallmark mitotic spindle and G2M cell cycle checkpoint pathways (**Supplementary Figure S2D**). Taken together, these data demonstrated that increased tumor aneuploidy correlates with CIN70 scores and cell proliferation pathways in CRCLM.

### CIN is linked to diminished adaptive immune responses

Recent evidence suggests that CIN may impact the anti-tumor immune response*(12)*. In this context, unsupervised hierarchical clustering of immune cell deconvolution patterns within the UCMC cohort identified two distinct immune clusters of CRCLMs (**Figure 3A**, clusters indicated by black and grey dendrogram colors). Metastases within the black cluster exhibited an “immune-enriched” gene expression pattern that is characteristic of cytotoxic cells, including natural killer (NK) and CD8+ T cells, B cells, and dendritic cells (DCs). By contrast, metastases within the grey cluster exhibited an “immune-depleted” gene expression pattern. Subtype 1 canonical metastases were highly enriched in the immune-depleted cluster (OR = 7.68, Chi squared p = 0.00018), whereas subtype 2 metastases were correspondingly enriched in the immune-enriched cluster (OR = 4.44, Chi squared p = 0.0017) (**Figure 3B**). A complete pair-wise comparison of the immune cell deconvolution patterns across the three integrated molecular subtypes of CRCLM confirmed that several immune cell populations, including B, T, NK, and DC cells, were decreased in canonical metastases (**Supplementary Figure S3A**).

**Figure 3.**
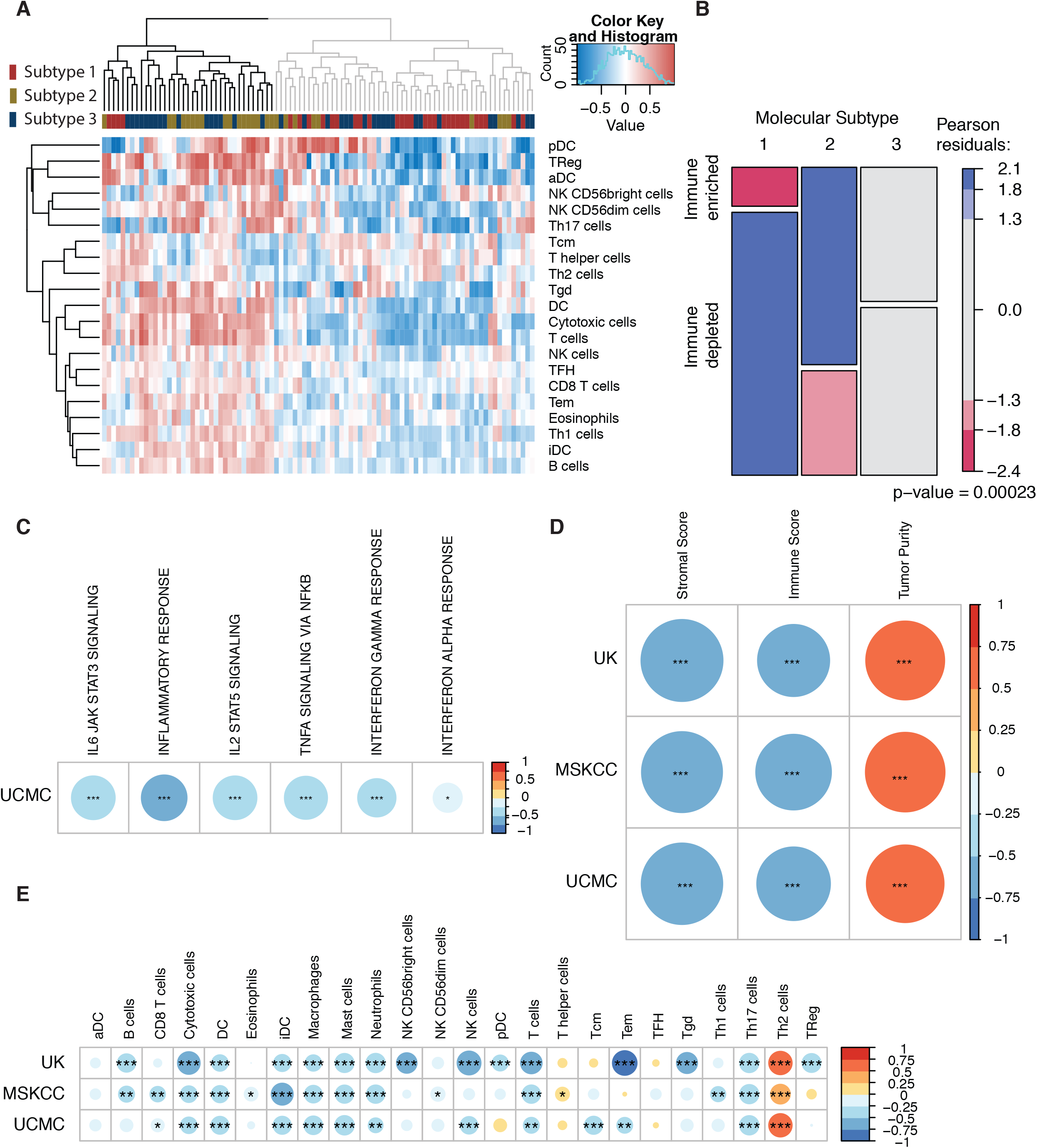
CIN is linked to diminished adaptive immune responses. **A)** Heatmap based on unsupervised hierarchical clustering showing the Bindea immune score signatures for each sample in the UCMC dataset; each row corresponds to an immune cell type, while each column corresponds to a patient sample; the horizontal bar above the heatmap indicates the integrated molecular subtype of the sample; the heatmap is color coded with the normalized Bindea scores with red, white, and blue indicating positive, zero, and negative values; unsupervised hierarchical clustering was performed on both rows and columns; the column dendrogram was color coded black for the immune-enriched group and grey for the immune-depleted group (n = 93). **B)** Mosaic plot showing the proportions of patients belonging to the immune-enriched and immune-depleted groups (defined by the black and grey clusters within the dendrogram shown in panel **A**) for each of the integrated molecular subtypes 1, 2, and 3; the rectangles are color-coded according to the Pearson residuals for the group, with blue corresponding to a positive residual or greater than expected, while red corresponds to a negative residual or less than expected; p-value was calculated using a chi-square test (n = 93). The height of the rectangular boxes corresponds to the proportion of the given molecular subtype in the immune-depleted or immune-enriched clusters. **C-E)** Correlation plots showing CIN70 correlated with different pathways or signatures; circle size and color are proportional to the Pearson correlation; red denotes a positive correlation, while blue indicates a negative correlation; *P < 0.05, **P < 0.01, and ***P < 0.001 (n = 93, 96, and 147 for UCMC, MSKCC, and UK/New-EPOC respectively). **C)** Correlation plot of CIN70 scores in the UCMC dataset with the ssGSEA scores for the Hallmark immune pathways. **D)** Correlation plot of the CIN70 scores in the three cohorts (UCMC, MSKCC, and UK/New-EPOC) with the ESTIMATE scores for the Stromal, Immune, and Tumor Purity signatures. **E)** Correlation plot of CIN70 scores in the three cohorts (UCMC, MSKCC, and UK/New-EPOC) with the normalized Bindea immune deconvolution scores.

We next examined whether there was an association between CIN70 scores and the expression of immunologic pathways of the Hallmark gene sets in the UCMC cohort of CRCLM. Indeed, a negative correlation was observed between CIN70 and ssGSEA scores for the immune pathways, including interferon alpha and gamma responses, IL6-JAK-STAT3 signaling, and TNF-alpha signaling (**Figure 3C**). We also applied the Estimation of STromal and Immune cells in MAlignant Tumours using Expression data (ESTIMATE) algorithm to estimate the stromal and immune composition of tumors*(13)*. We found that CIN70 was positively correlated with tumor purity, but negatively correlated with stromal and immune composition in all three cohorts (Pearson correlation p < 0.0001) (**Figure 3D**). These findings suggested that high CIN is associated with diminished expression of immunologic pathways and immune composition within CRCLMs.

We further characterized the tumor microenvironment within high CIN70 metastases using three distinct immune cell deconvolution analyses. In all three CRCLM cohorts, we observed an inverse correlation between CIN70 scores and signatures indicative of adaptive cytolytic immune cells (**Figure 3E** and **Supplementary Figure S3B**). Similarly, in the UK/New-EPOC cohort the Microenvironment Cell Populations-counter (MCP-counter) method*(14)* identified an inverse correlation between CIN70 scores and the abundance of immune cell populations, including B, CD8+ T, cytotoxic, NK, and dendritic cells (**Supplementary Figure S4**). Collectively, these data demonstrated that elevated CIN70 is associated with low intratumoral infiltration of adaptive immune cells in CRCLM.

### CIN is associated with sensitivity to DNA-damaging agents

We previously reported that a subset of patients with canonical metastases experienced favorable overall survival rates similar to the immune subtype of CRCLM, while most experienced inferior outcomes*(1)*. Within the cohort of canonical metastases, low CIN70 was associated with improved overall survival when compared to high CIN70 CRCLM (HR = 0.42 for low vs. high CIN70, univariate Cox p = 0.0041). We hypothesized that the improved clinical outcomes of these patients are related to increased chemosensitivity of high CIN metastases to DNA-damaging therapies, such as topoisomerase inhibitors, which are commonly utilized in the treatment of colorectal cancers. We examined the relationship between CIN70 values and sensitivity to topotecan and irinotecan in 246 carcinoma cell lines from the Cancer Cell Line Encyclopedia (CCLE) (**Figure 4A**). Drug sensitivity was determined over a range of drug doses and reported as the IC50 value for each cell line. We found that the carcinoma cell lines with the highest levels of CIN70 displayed increased sensitivity (lowest IC50 values) to topotecan (one-way ANOVA FDR-corrected p < 0.0001) and irinotecan (one-way ANOVA FDR-corrected p < 0.0001). Similarly, high CIN70 cell lines were also sensitive to other DNA damaging therapies, including ionizing radiation (one-way ANOVA p = 0.014). By contrast, we observed no relationship between CIN70 and increased sensitivity to agents not considered to elicit DNA damage, such as paclitaxel (one-way ANOVA FDR-corrected p = 0.38). Indeed, a complete analysis of all drugs tested in the CCLE showed that only topotecan and irinotecan had a statistically significant correlation with CIN70 (**Supplementary Figure S5A**). We examined an additional set of 4,686 drug compounds available in the DepMap database in which cell viability was measured after treatment with a single drug concentration. The results showed that cell viability after treatment with topoisomerase inhibitors exhibited the greatest negative correlation with CIN70 score (**Supplementary Figure S5B)**. We also found an inverse relationship between CIN70 scores and cell viability after oxaliplatin treatment, which is commonly utilized in the treatment of CRCLM (**Supplementary Figure S5C**). Taken together, these findings demonstrated that high CIN70 scores are associated with increased sensitivities to DNA-damaging cancer therapies.

**Figure 4.**
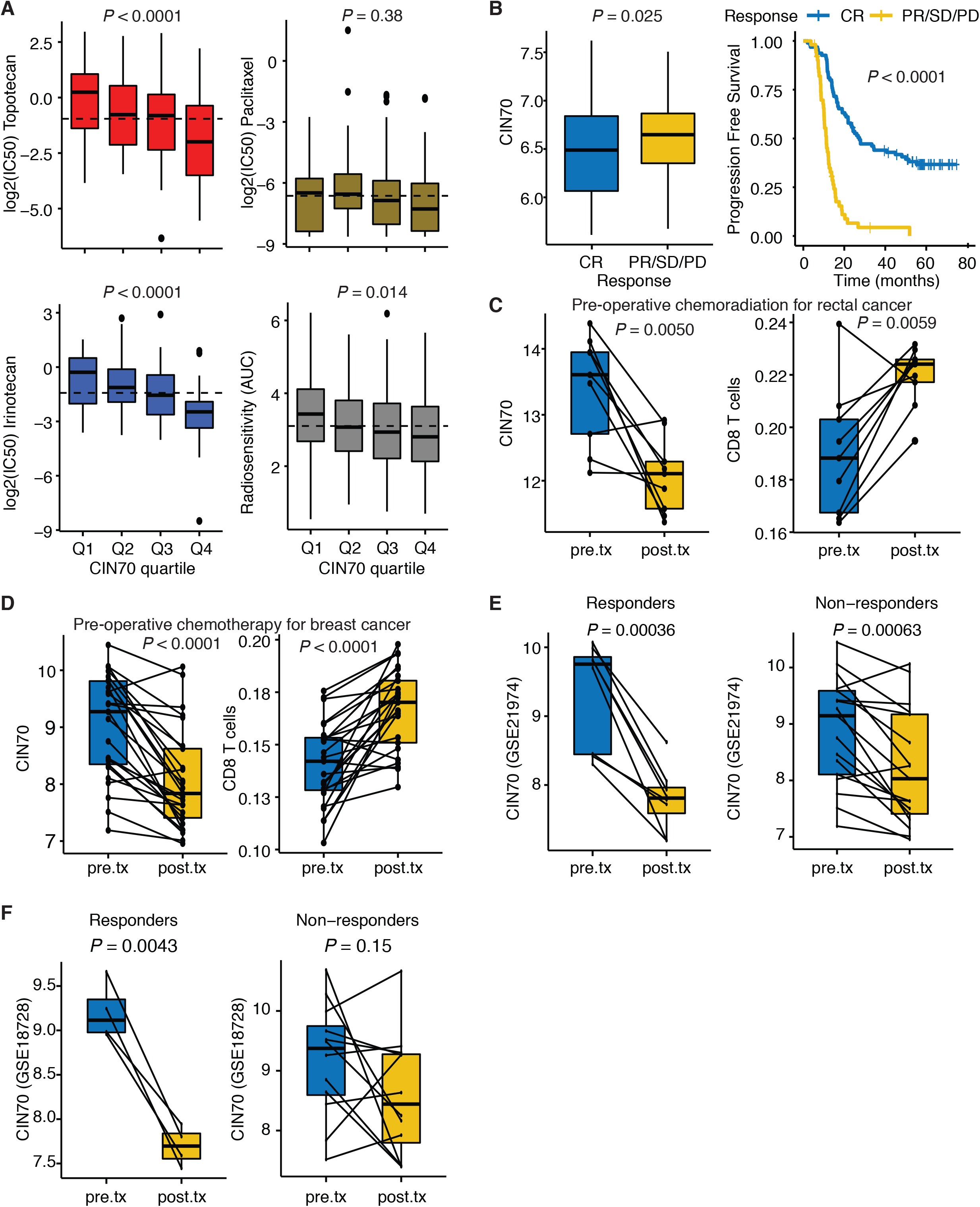
DNA-damaging agents decrease CIN and augment adaptive immunity. **A)** Top left, top right, and bottom left panels show the log(IC50) of CCLE cell lines versus CIN70 quartiles for topotecan (red, n = 222), irinotecan (blue, n = 141), and paclitaxel (olive green, n = 223), respectively; the bottom right panel shows a boxplot of radiosensitivity of CCLE cell lines measured as area under the curve (AUC) for different CIN70 quartiles (n = 455); p-values were calculated using one-way ANOVA; p-values for topotecan, irinotecan, and paclitaxel were derived from a common CCLE drug screen and corrected for multiple comparisons using the Benjamini-Hochberg method for false discovery rate (see **Supplementary Figure 5**). **B)** Boxplot showing CIN70 scores for patient samples in the UK/New-EPOC dataset who had a complete response (CR) or partial response (PR)/stable disease (SD)/progressive disease (PD) based on RECIST principles using CT imaging following pre-operative chemotherapy (left panel); p-value was calculated using a two-tailed t-test. PR/SD/PD tumors were grouped together since there was no difference in CIN70 scores among these three response categories (one-way ANOVA p=0.99). Kaplan-Meier curves of the UK/New-EPOC dataset for progression free survival based on response to chemotherapy (right panel); p-value for statistical difference was calculated using the log-rank method (n = 147). **C)** Paired boxplots showing CIN70 score (left panel) and CD8+ T cell score (right panel) changes between pre-treatment (blue) and chemoradiation post-treatment samples (yellow) of rectal cancer dataset GSE15781 (n = 9); p-values were calculated using a two-tailed paired t-test. **D)** Paired boxplots showing CIN70 score (left panel) and CD8+ T cell score (right panel) changes between pre-treatment (blue) and chemotherapy post-treatment samples (yellow) of breast cancer dataset GSE21974 (n = 25); p-values were calculated using a two-tailed paired t-test. **E)** Paired boxplots showing CIN70 score changes between pre-treatment (blue) and post-treatment (yellow) samples in tumors that showed a pathologic complete response (responder) (left panel, n = 8) and in tumors that did not show a pathologic complete response (non-responder) (right panel, n = 17) in breast cancer dataset GSE21974; p-values were calculated using a two-tailed paired t-test. **F)** Paired boxplots showing CIN70 score changes between pre-treatment (blue) and post-treatment (yellow) samples in responding tumors (left panel, n = 4) and non-responding tumors (right panel, n = 12) in breast cancer dataset GSE18728; p-values were calculated using a two-tailed paired t-test.

### DNA-damaging agents deplete high-CIN tumor cells and augment adaptive immunity

In the UK/New-EPOC randomized clinical trial, wherein CRCLM patients received treatment with pre-operative oxaliplatin/irinotecan with 5-FU/capecitabine, we found that patients whose tumors exhibited a complete radiographic response (CR) to pre-operative chemotherapy versus patients whose tumors demonstrated a partial response (PR), stable disease (SD), or progressive disease (PD) harbored a lower post-chemotherapy CIN70 score (mean CIN70 = 6.46 for CR vs. 6.62 for PR/SD/PD, two-tailed t-test p = 0.025) (**Figure 4B**, left panel) in association with markedly improved progression-free survival (HR = 0.22 [0.14 – 0.33, 95% CI], univariate Cox p < 0.0001) (**Figure 4B**, right panel). Since high CIN70 is associated with sensitivity to DNA-damaging agents and diminished anti-tumor immune responses, we investigated whether treatment with DNA-damaging drugs might deplete high-CIN populations of cells within tumors and thereby improve host adaptive immunity.

We tested this hypothesis in four additional independent clinical cohorts. First, in patients with primary rectal cancer receiving pre-operative chemoradiation therapy with fractionated radiation and concurrent capecitabine chemotherapy (GSE15781), we compared matched pre-treatment (before) and post-treatment (4-6 weeks after chemoradiation therapy) tumor specimens. This analysis demonstrated lower CIN70 scores (two-tailed paired t-test p = 0.0050, mean = 13.40 before vs. 12.07 after) and higher CD8+ T cell infiltration (two-tailed paired t-test p = 0.0059, mean = 0.19 before vs. 0.22 after) in post-treatment tumors as compared to matched pre-treatment samples (**Figure 4C** and **Supplementary Figure S6A**). Consistent with these findings, in a second cohort of matched pre-treatment and post-treatment breast cancer biopsies from women receiving pre-operative epirubicin, cyclophosphamide, and docetaxel chemotherapy (GSE21974), CIN70 decreased in post-treatment samples (two-tailed paired t-test p < 0.0001, mean = 9.02 before vs. 8.08 after), which was associated with a concomitant increase in tumor CD8+ T cells (two-tailed paired t-test p < 0.0001, mean = 0.14 before vs. 0.17 after) (**Figure 4D**, left and right panels; **Supplementary Figure S6B**). Likewise, a third cohort of matched pre-treatment and post-treatment breast cancer biopsies from women receiving pre-operative capecitabine and docetaxel chemotherapy (GSE18728) showed a decrease in CIN70 following treatment (two-tailed paired t-test p = 0.016, mean = 9.19 before vs. 8.37 after), as well as an increase in T cell signature (two-tailed paired t-test p = 0.0020, mean = 0.017 before vs. 0.10 after) (**Supplementary Figure S7A** and **S7B)**. CIN70 scores were greatest in triple-negative breast cancers relative to ER/PR+ and HER2+ breast cancers (**Supplementary Figure S7C**) consistent with evidence for increased chromosomal instability in the triple-negative subtype of breast cancers*(15)*. However, the change in CIN70 during neoadjuvant chemotherapy was observed across all subtypes of breast cancer (**Supplementary Figure S7D**). Similar results were observed in a fourth cohort consisting of patients with muscle-invasive bladder cancer treated with neoadjuvant methotrexate, vinblastine, doxorubicin, and cisplatin chemotherapy prior to radical cystectomy (**Supplementary Figures S8A** and **S8B**).

Importantly, in women receiving pre-operative chemotherapy for breast cancer, pathologic complete responders (pCR) demonstrated a larger decrease in CIN70 than pathologic non-responders (non-pCR). The difference in CIN70 scores between matched pre-treatment and post-treatment samples for women receiving pre-operative epirubicin, cyclophosphamide, and docetaxel chemotherapy (GSE21974) was 1.53 for pCR (mean = 9.32 before vs. 7.79 after, two-tailed paired t-test p = 0.00036) and 0.67 for non-pCR tumors (mean = 8.88 before vs. 8.21 after, two-tailed paired t-test p = 0.00063), respectively. Similarly, the mean difference of CIN70 scores in women receiving capecitabine and docetaxel chemotherapy (GSE18728) was 1.52 for pCR (two-tailed paired t-test p = 0.0043) and 0.59 non-pCR tumors (two-tailed paired t-test p = 0.15), respectively (**Figure 4E and 4F**, left and right panels). Taken together, these data demonstrated that elevated CIN70 scores are associated with increased sensitivity to DNA-damaging therapies, which attenuate CIN and increase intratumoral immune responses in human cancers. Importantly, the magnitude of decrease in CIN70 is associated with pathologic complete response to pre-operative chemotherapy treatment.

## Discussion

In this study, we demonstrated that the canonical subtype of CRCLM is enriched for cells with high levels of CIN. We also found that the CIN70 expression signature serves as a reliable surrogate for CIN, wherein the lowest quartile of CIN70 scores associated with low levels of aneuploidy. We propose that CIN70 might be utilized as a clinical biomarker that is complementary to other existing biomarkers, including signatures of cell proliferation*(16)*. Furthermore, we found that high CIN70 tumors exhibit increased metastatic potential, which in turn portends adverse progression-free and overall survival rates among patients with CRCLM. These poor patient outcomes may be caused by low immune responses that we observed in canonical CRCLM tumors with high CIN70 scores. By contrast, however, these CIN-rich tumors are particularly vulnerable to DNA-damaging therapies, such as topoisomerase inhibitors. Taken together, our results demonstrate how CIN70 may help guide therapeutic decision making in CRCLM.

Thus far, it remains unclear how CIN affects cancer cell fitness. Some studies suggest that CIN promotes immune evasion and mesenchymal phenotypes which can in turn enable metastasis, particularly in sub-optimal growth conditions, such as under selective pressure from cytotoxic agents or the loss of an oncogenic mutation*(17, 18)*. Furthermore, metastatic tumors exhibit more aneuploidy than their localized counterparts, an observation which could be partially explained through a mechanism of increased phenotypic plasticity*(19)*. However, excessively high levels of chromosomal instability in cancer cells can be cytotoxic and thereby reduce cellular proliferation*(17, 20)*, potentially mediated via innate immune signaling through the cGAS-STING pathway*(21)*. Regardless of the CIN70 score, most tumors in our cohort did not exceed an aneuploidy score of approximately 0.5, suggesting that unbalanced copy number aberrations involving more than half the genome may be cytotoxic. Therefore, there may exist an optimal level of CIN to support cancer cell fitness that enables aggressive phenotypes without compromising cell viability. In our cohort, CIN70 displays a non-linear relationship with aneuploidy scores and various endpoints, suggesting different thresholds for specific malignant phenotypes. In the case of metastasis, tumors with the greatest CIN70 scores showed the greatest predilection for metastasis to high-risk/adverse prognosis organ sites.

Immune signaling may mediate the paradoxical relationships between CIN and tumor fitness. We observed that various types of immune cell, including B cells, T cells, NK cells, and professional antigen presenting cells, are depleted in high-CIN CRCLMs. Other studies have demonstrated similar findings, wherein higher levels of aSCNAs are associated with markers of immune evasion, including a reduction in tumor-infiltrating CD8+ T cells*(22)*. Similarly, the downregulation of type I interferon signaling and upregulation of the NF-κB are known to protect high-CIN tumors from inflammation*(9)*. Furthermore, aSCNAs that evolve during treatment may lead to the disappearance of neoantigens through loss of heterozygosity encompassing these genomic loci*(23)*.

Despite the apparent inverse relationship between CIN70 and immune pathways, many of the aberrant karyotypes and frameshift mutations generated by CIN can serve to function as immunogenic epitopes, potentially enhancing immune surveillance of cancer cells*(20, 24)*. Polyploid tumors can also trigger immunogenic mechanisms through endoplasmic reticulum-associated stress where calreticulin is exported from the cell and recognized by CD8+ T-cells and NK cells*(23)*. It appears then, that CIN plays a multifactorial role in tumorigenesis and progression, wherein some degree of CIN supports oncogenesis, while levels above a certain threshold may trigger the adaptive immune response that promotes cancer cell death.

We also found that high-CIN tumors exhibit sensitivity to DNA-damaging therapies, which presents an opportunity to improve patient outcomes. Specifically, we find that topoisomerase inhibitors are highly effective against cancer cells with high CIN70. This observation is consistent with prior reports showing that chromosomal untangling and packaging is defective in high CIN cancer cells*(25)*. Because of this sensitivity, chemotherapy appears to deplete high-CIN70 cells from primary rectal, breast, and bladder cancers, resulting in improved anti-tumor immune responses and favorable clinical outcomes. Taken together, we propose that CIN represents a powerful predictive biomarker to guide treatment decision making.

In summary, we found that CIN plays a central and complex role in CRCLM, wherein genomic instability simultaneously promotes protective and deleterious effects on tumor fitness. High CIN tumors exhibit elevated metastatic potential, leading to unfavorable baseline prognoses for patients with CRCLM. However, these same genomic aberrations reveal a vulnerability to DNA-damaging therapies, thereby presenting an opportunity to personalize treatments for these high-risk patients. Importantly, we also demonstrated that our observations are not limited to colorectal cancer; similar results were observed across several cancer types in our study, including breast and bladder cancer. Further studies are needed to explore the breadth of the applications of CIN to optimize patient outcomes through selecting therapies to which a given tumor is likely most sensitive. Thus, it is important to validate these observations in prospective clinical trials to confirm whether CIN may serve as an important clinical biomarker of response to genotoxic chemotherapy and provide tailored treatments to patients across cancer types.

## Materials and Methods

### Data sources

The normalized gene expression and drug sensitivity data (log2 (IC50)) were downloaded from the Cancer Cell Line Encyclopedia (CCLE) website (https://portals.broadinstitute.org/ccle) and the DepMap Portal (https://depmap.org/portal/). Radiosensitivity data, defined by the area under the curve (AUC), were obtained from Yard et al*(26)*. Normalized gene expression data for GSE21974, GSE18728, GSE15781, and GSE48277 were downloaded from the GEO data repository (https://www.ncbi.nlm.nih.gov/geo/). The UCMC dataset was previously described and is available for download from the European Genome-Phenome Archive (https://ega-archive.org/studies/EGAS00001002945). The MSKCC dataset was downloaded from the ArrayExpress website (https://www.ebi.ac.uk/arrayexpress/experiments/E-MTAB-1951/). The UK/New-EPOC dataset was downloaded from a privately accessed cBioPortal request following MTA approval by the Stratification in Colorectal Cancer (S:CORT) consortium. The MSK-IMPACT dataset was downloaded from the Metastatic Colorectal Cancer (MSKCC, Cancer Cell 2018) study on cBioPortal*(27, 28)*. Progression-free survival was defined as the interval between the start of neoadjuvant chemotherapy and disease progression or death (event) or last follow-up (censor). Disease-free survival was defined as the interval between surgical resection of all visible disease and disease progression or death (event) or last follow-up (censor). Overall survival was defined as the interval between the end of treatment and death (event) or last follow-up (censor).

Patients with CRC liver metastases received therapies that were considered standard-of-care at the time of treatment. In the UCMC, MSKCC, and UK/New-EPOC cohorts, patients predominantly received perioperative systemic therapy, consisting of 5-fluorouracil-based chemotherapy typically combined with oxaliplatin and/or irinotecan, curative intent management of primary colorectal tumors, and partial hepatectomy of all visible liver metastases. In the UK/New-EPOC cohort, ~50% of the patients also received peri-operative cetuximab. 221 of 257 patients in the UK/New-EPOC cohort underwent surgery after peri-operative chemotherapy of which 147 CRC liver metastases successfully underwent molecular analysis.

### Microarray analysis

CEL files from the UK/New-EPOC dataset were downloaded and processed using the Affymetrix Array Power Tools (APT) (https://www.thermofisher.com/us/en/home/life-science/microarray-analysis/microarray-analysis-partners-programs/affymetrix-developers-network/affymetrix-power-tools.html). Microarray data were background and RMA normalized prior to analysis.

### RNA sequencing analysis

RNA-seq analysis was carried out by first aligning 75 bp length paired-end reads to the hg38 human reference genome using the *STAR* aligner version 2.6.1d*(29)*. The resulting .bam files were then sorted using *samtools* version 1.10*(30)*. The total reads per gene were counted using *htseq* in stranded mode with the Ensembl hg38 list of coding exons for each gene as a reference. A matrix of counts for each gene and sample was then generated. The R package *edgeR(31)* was used to generate a table of logCPM values for each gene.

### CIN70 score calculation

CIN70 score calculation was performed as previously described*(7)*. In brief, the average of the normalized gene expression data of the 70 genes that constitute the signature was used to calculate the CIN70 score in each dataset. In the case of microarray datasets such as GSE21974, GS18728, GSE48277, GSE15781, MSKCC, and UK/New-EPOC, the probe with the maximum RMA-normalized average signal for each gene in the signature was used for calculating CIN70. For RNA-seq data such as the UCMC dataset, the CIN70 signature was calculated using the *ssgsea* method implemented in the R package EGSEA*(32)* with a custom gene signature containing the 70 genes.

### Recurrent copy-number variation detection

GISTIC 2.0*(33)* v6.15.28 was run using the GenePattern*(34)* platform to identify significantly recurrent regions of copy-number variation separately in the UCMC cohort and the *Yaeger et al* MSKCC validation cohort*(27)*. A pseudomarker file was used for both cohorts with the default spacing value of 10,000 bases.

### Arm-level somatic copy number alteration calls and aneuploidy score

Arm-level somatic copy number alterations (aSCNAs) were called using ASCETS*(10)* with the default parameters (log_2_ copy ratio threshold = ±0.2; arm alteration fraction threshold = 0.7; min breadth of coverage [BOC] = 0.5). Aneuploidy scores were calculated as previously described*(35)* for each sample by computing the number of arms affected by aSCNAs (ASCETS call = AMP or DEL). However, panel data can result in insufficient BOC to make reliable aSCNA calls on some chromosome arms (ASCETS call = LOWCOV). Thus, we modified the score by dividing the number of arms with aSCNAs by the total number of arms with sufficient BOC in the sample (ASCETS call = AMP, DEL, NEUTRAL, or NC). The final scores represent the fraction of evaluable arms harboring aSCNAs in each sample.

### Immune deconvolution

Immune deconvolution analysis was carried out by using the xCell*(36)* webtool (https://xcell.ucsf.edu). For the UCMC RNA-seq dataset the input data was the FPKM values for each gene in the sample. For the GSE21974, GS18728, GSE48277, GSE15781, MSKCC and UK/New-EPOC microarray datasets, the probe level data was converted to gene symbols and the xCell tool was ran with the RNA-seq data option unselected.

### ESTIMATE analysis and MCP counter score

The Estimation of STromal and Immune cells in MAlignant Tumours using Expression data (ESTIMATE) algorithm*(13)* was implemented using the R “estimate” package (https://bioinformatics.mdanderson.org/public-software/estimate/). The logCPM data was used for the analysis of the UCMC dataset. Similarly, the MCP counter score was carried out using the “MCPcounter” package in R (https://cit.ligue-cancer.net/mcp-counter/).

### Statistical analysis

Analyses were performed using R version 3.5.1. Data were analyzed by one-way analysis of variance (ANOVA), two-tailed Student’s t test, log-rank test, and chi-square tests. A Shapiro-Wilk test was used to test for normality when appropriate. Kaplan-Meier curves were generated using the survminer package in R; a log-rank test was used to compare survival between groups. Hazard ratios were calculated using a Cox proportional hazards regression model. Meta-analysis of hazard ratios was carried out using the *rmeta* package in R. In the case of matched data, a paired two-tailed Student’s t-test was used. Correlation analysis was done by calculating the Pearson correlation between the variables; the *corrplot* R package was used for generating the correlation plots. P-values were corrected for multiple comparisons using the Benjamini-Hochberg method for false discovery rate. Statistical significance was denoted according to the following: *P < 0.05, **P < 0.01, ***P < 0.001, ****P < 0.0001.

## Acknowledgments

This work was supported by the Ludwig Cancer Research Foundation (SPP and RRW).

## Author Contributions

Conceptualization: SPP

Methodology: CAM, LFS, SPP

Data acquisition: SAP, JAB, JNP, ED, TSM, MID, SPP

Data analysis: CAM, LFS, SPP

Visualization: CAM, LFS, SPP

Funding acquisition: SPP

Supervision: SPP

Writing – original draft: CAM, LFS, SCI, SPP

Writing – review & editing: All authors

## Competing Interests

SPP and RRW are co-inventors on US patents titled “Methods and Kits for Diagnosis and Triage of Patients with Colorectal Liver Metastases” and “Molecular Subtyping of Colorectal Liver Metastases to Personalize Treatment Approaches”. All other authors have no conflicts to disclose.

## Data and Materials Availability

All data and code were obtained from existing public sources with the following exceptions: (1) UK/New-EPOC genomic and clinical data required MTA approval from S:CORT consortium and (2) clinical data for MSKCC cohort required MTA approval from MSKCC.

## Supplementary Figure Legends

**Supplementary Figure S1.**
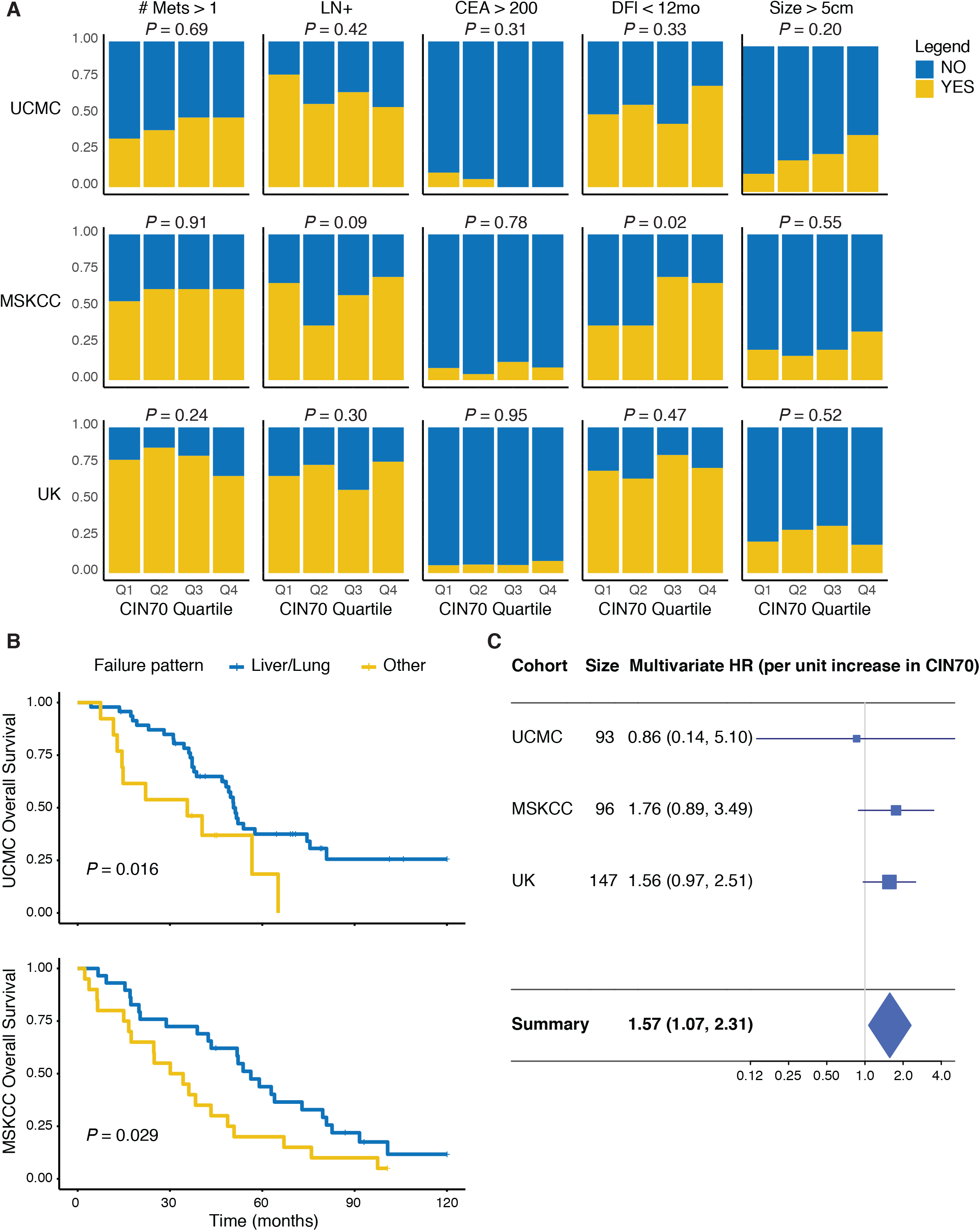
Risk association of patient cohorts with CIN70 and patterns of failure. **A)** Stacked barplots showing the associations of Clinical Risk Score (CRS) factors with CIN70 quartiles for the UCMC, MSKCC, and UK/New-EPOC datasets; yellow denotes the proportion of patients that exhibit each factor, while blue denotes the proportion of patients that do not; p-values were calculated using a chi-square test (n = 93, 96, and 147 for UCMC, MSKCC, and UK/New-EPOC respectively). LN+, lymph node positive. CEA, carcinoembryonic antigen. DFI, disease-free interval between primary tumor and presentation of liver metastasis. **B)** Kaplan-Meier curves showing overall survival for patients with liver/lung metastasis (blue) or metastasis to “other” sites (yellow); top panel corresponds to the UCMC dataset (n = 60 patients with recurrence data) and bottom panel corresponds to the MSKCC dataset (n = 49 patients with recurrence data); p-values were calculated using a log-rank test. **C)** Meta-analysis for the multivariate hazard ratio of overall survival based on CIN70 (continuous variable) and Clinical Risk Score (dichotomized at <2 or >=2); horizontal blue lines denote the 95% CI; standard errors were calculating assuming random effects; horizontal axis shows the hazard ratio on a log scale; the boxes show the CIN70 effect estimates from the single studies, while the diamond shows the pooled result (n = 93, 96, and 147 for UCMC, MSKCC, and UK/New-EPOC respectively).

**Supplementary Figure S2.**
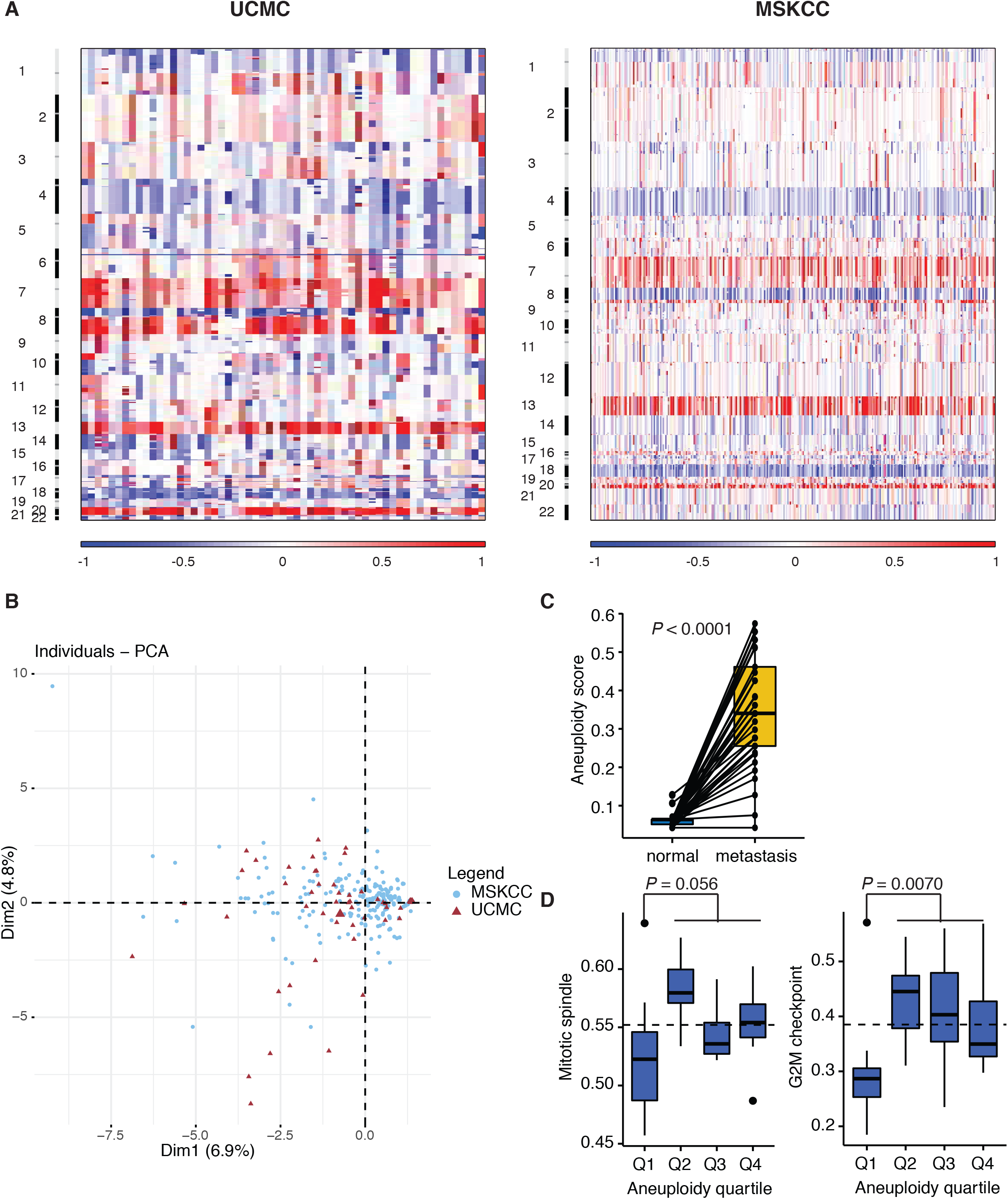
UCMC and MSKCC cohort samples show similar pattern of amplifications and deletions. **A)** Heatmap plots showing the overall copy-number profiles of the UCMC cohort (left, n = 59) and MSKCC cohort (right, n = 315) generated by the GISTIC 2.0 algorithm; columns correspond to patient samples, while rows correspond to chromosomal loci; red indicates amplification, while blue indicates deletion. **B)** PCA plot showing the distribution of the UCMC (red triangle, n = 59) and MSKCC (blue circle, n = 315) cohorts; horizontal axis denotes the dimension that explains the largest variance, while the vertical axis denotes the dimension that explains the second largest variance. **C)** Paired plot showing the aneuploidy scores in matched normal liver and CRC liver metastasis samples in UCMC dataset (n = 45); p-value was calculated using a two-tailed paired t-test. **D)** Boxplots showing the mitotic spindle and G2M checkpoint ssgsea scores as a function of the aneuploidy score quartile in the UCMC dataset (n = 32); p-values were calculated using one-way ANOVA.

**Supplementary Figure S3.**
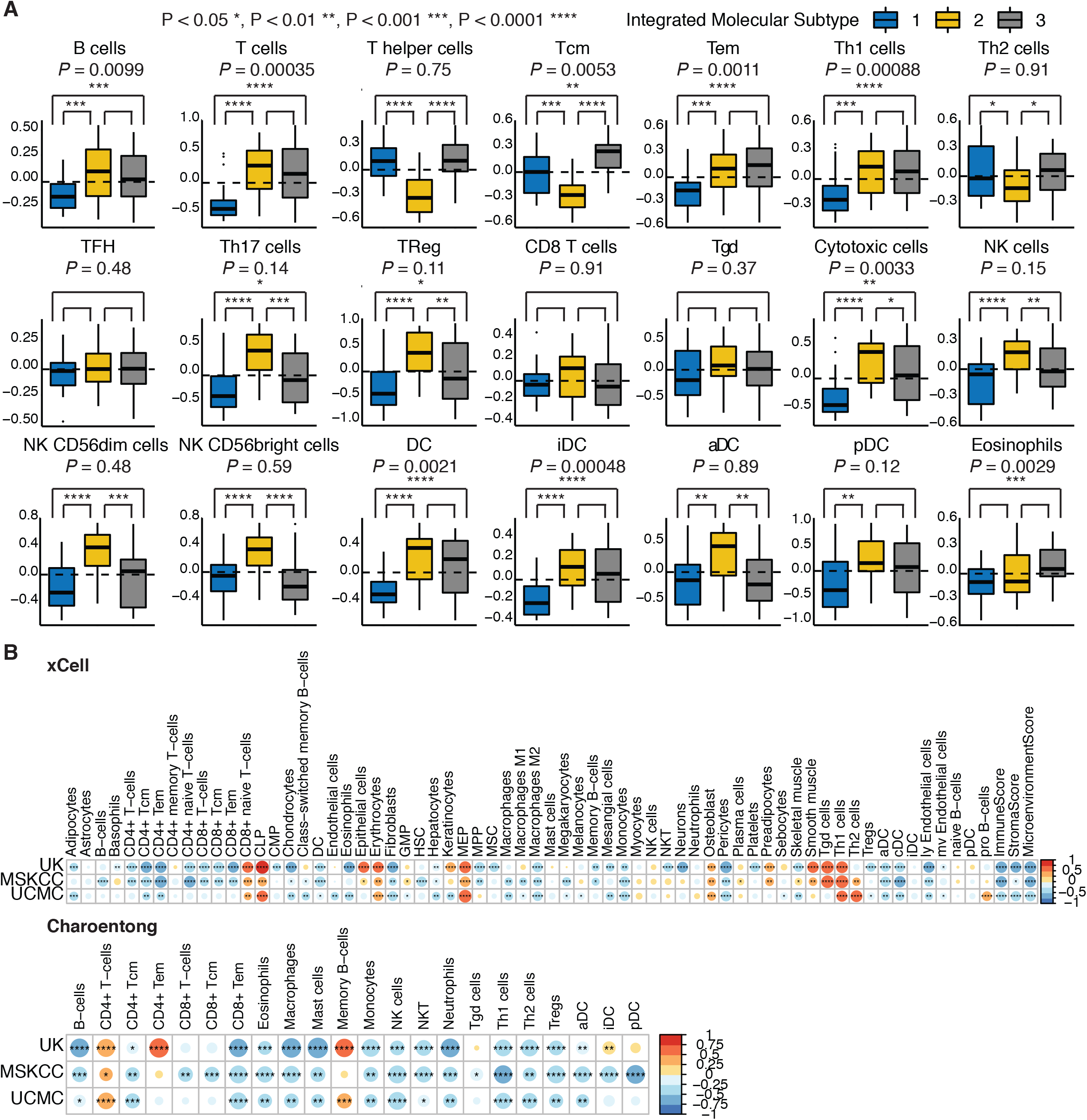
Immune deconvolution analysis of patient cohorts. **A)** Boxplots showing the specific cell type Bindea immune deconvolution scores for the three patient subtypes in the UCMC dataset (n = 93); integrated molecular subtypes are indicated by the colors blue, yellow, and grey, respectively; p-values above subplots were calculated using a one-way ANOVA test; paired statistical analysis was calculated using a two-tailed t-test; p-values were corrected for multiple comparisons using the Benjamini-Hochberg method for false discovery rate. **B)** Pearson correlation plots of the xCell (top panel) and Charoentong (bottom panel) immune deconvolution cell type signatures and CIN70 score for the UK/New-EPOC, MSKCC, and UCMC cohorts; circle size and color are proportional to the Pearson correlation; blue denotes a negative correlation, while red denotes a positive correlation; *P < 0.05, **P < 0.01, ***P < 0.001, and ****P < 0.0001 (n = 93, 96, and 147 for UCMC, MSKCC, and UK/New-EPOC respectively).

**Supplementary Figure S4.**
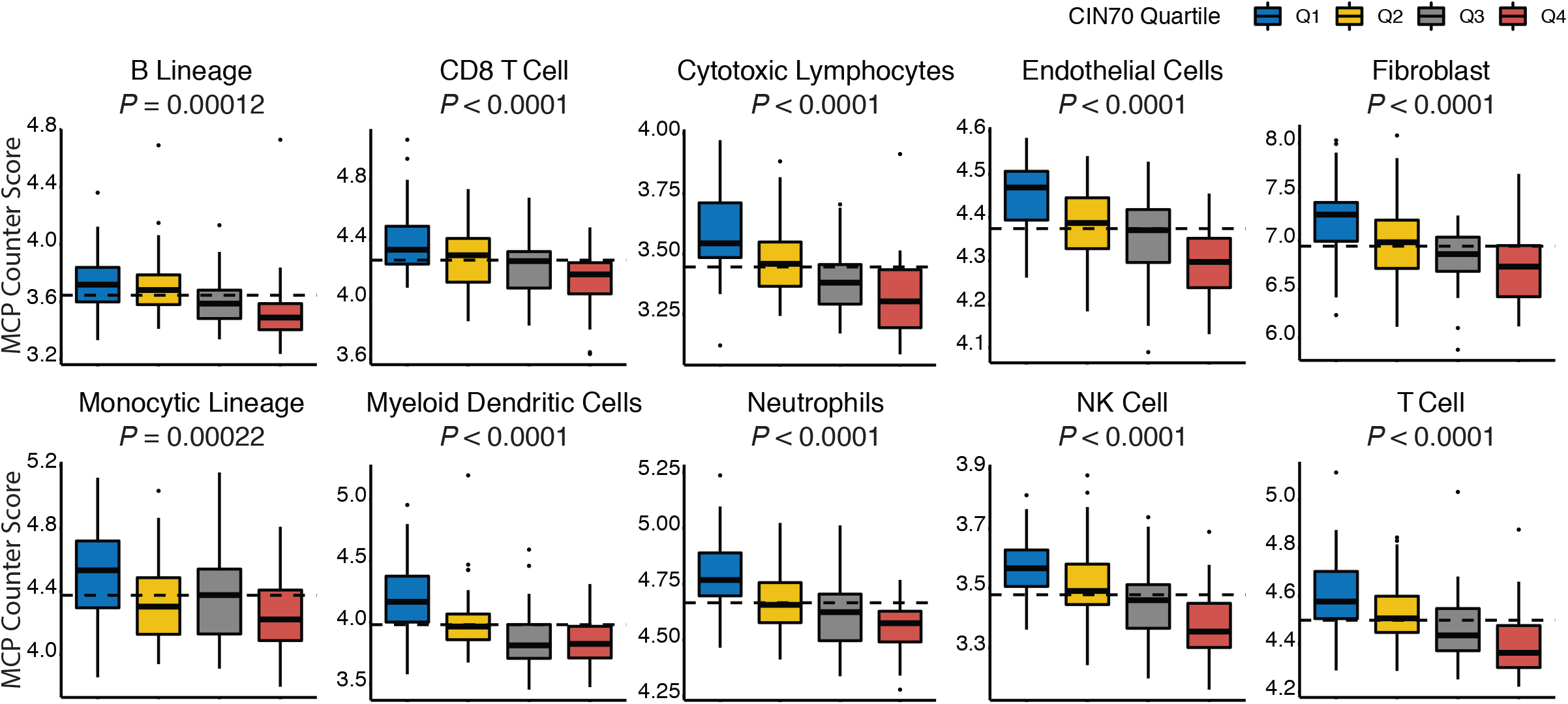
MCP Counter immune signatures negatively correlated with CIN70. Boxplots showing the MCP counter score of different cell types versus the CIN70 score quartiles in the UK/New-EPOC dataset (n = 147); p-value was calculated using a one-way ANOVA; p-values were corrected for multiple comparisons using the Benjamini-Hochberg method for false discovery rate.

**Supplementary Figure S5.**
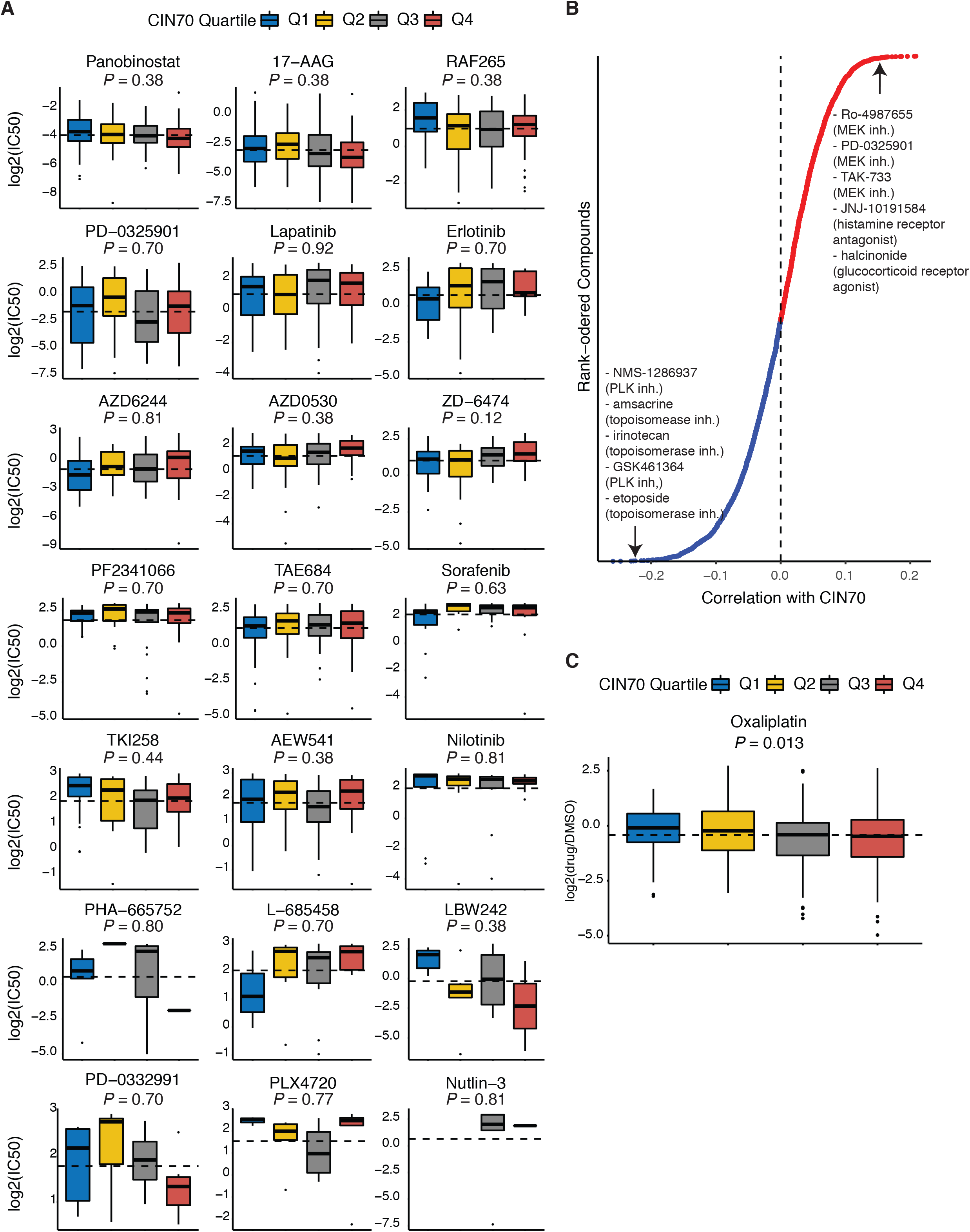
High CIN70 is associated with increased sensitivity to DNA-damaging compounds. **A)** Boxplots showing the log2 (IC50) of 21 chemotherapy compounds tested on 246 CCLE cell lines versus CIN70 quartiles; p-values were calculated using a one-way ANOVA; p-values were corrected for multiple comparisons using the Benjamini-Hochberg method for false discovery rate. **B)** Pearson correlation plot of cell viability and CIN70 for 4,686 drug compounds and 578 CCLE cell lines; red color indicates a positive correlation, while blue color indicates a negative correlation; the top five drug compounds at either end of the spectrum are listed along with the mechanism of action. **C)** Boxplot showing the log2 cell viability following treatment with oxaliplatin tested on 343 CCLE cell lines versus CIN70 quartiles; p-value was calculated using a one-way ANOVA.

**Supplementary Figure S6.**
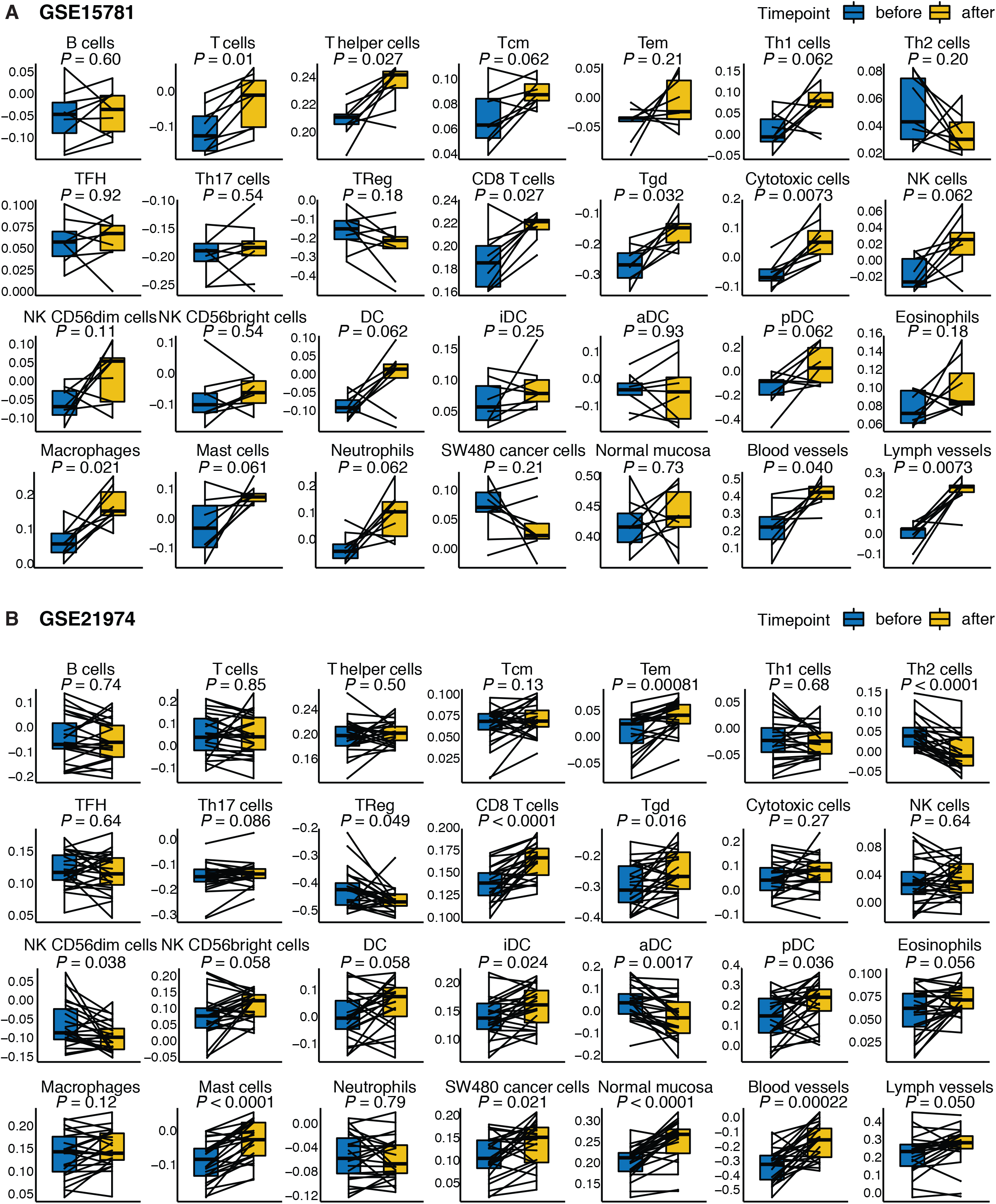
Immune cell signature changes during cancer treatment. **A-B)** Paired boxplots showing the changes in Bindea immune deconvolution scores during pre-operative chemoradiation therapy (fractionated radiation and concurrent capecitabine chemotherapy) for the rectal cancer dataset GSE15781 (n = 9) (**A**) and during neoadjuvant pre-operative chemotherapy (epirubicin, cyclophosphamide, and docetaxel chemotherapy) for the breast cancer dataset GSE21974 (n = 25) (**B**); p-values were calculated using a two-tailed paired t-test; p-values were corrected for multiple comparisons using the Benjamini-Hochberg method for false discovery rate.

**Supplementary Figure S7.**
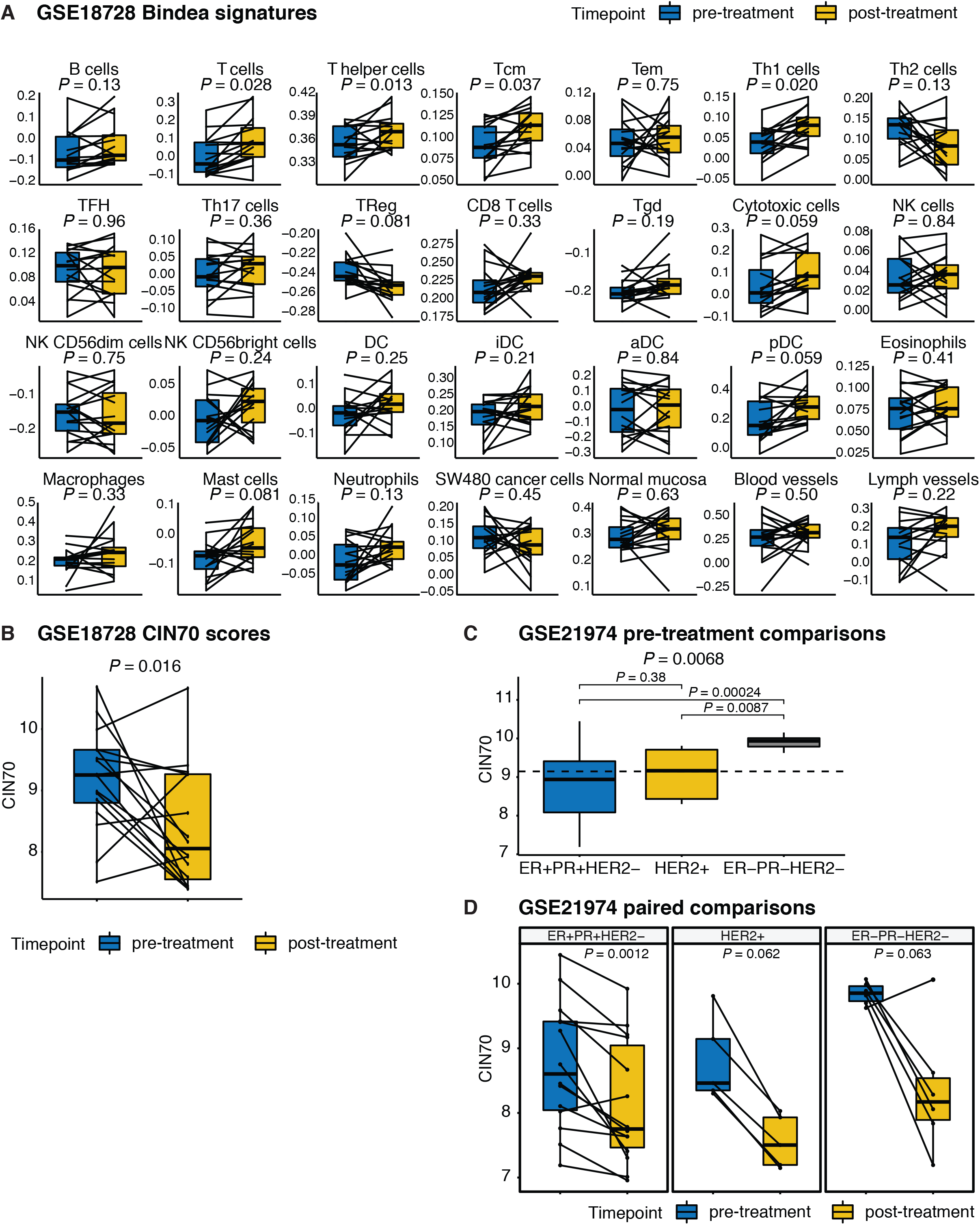
Immune cell signature and CIN70 score changes during neoadjuvant breast cancer treatment. **A)** Paired boxplots showing the Bindea immune deconvolution scores comparing the pre-treatment samples (blue) to the post-treatment samples (yellow) for breast cancer dataset GSE18728 (n = 16); p-values were calculated using a paired t-test; p-values were corrected for multiple comparisons using the Benjamini-Hochberg method for false discovery rate. **B)** Paired boxplot showing the change in CIN70 scores comparing the pre-treatment samples (blue) to the post-treatment samples (yellow) for breast cancer dataset GSE18728 (n = 16); p-value was calculated using a two-tailed paired t-test. **C)** Boxplots of CIN70 scores by breast cancer molecular subtype (based on ER, PR, and HER2 status) in GSE21974; p-values above subplots were calculated using a one-way ANOVA test; paired statistical analysis was calculated using a two-tailed t-test. **D)** Paired boxplots showing the change in CIN70 scores comparing the pre-treatment samples (blue) to the post-treatment samples (yellow) for breast cancer dataset GSE21974 (n = 25) as a function of molecular subtype; p-values were calculated using a two-tailed paired t-test.

**Supplementary Figure S8.**
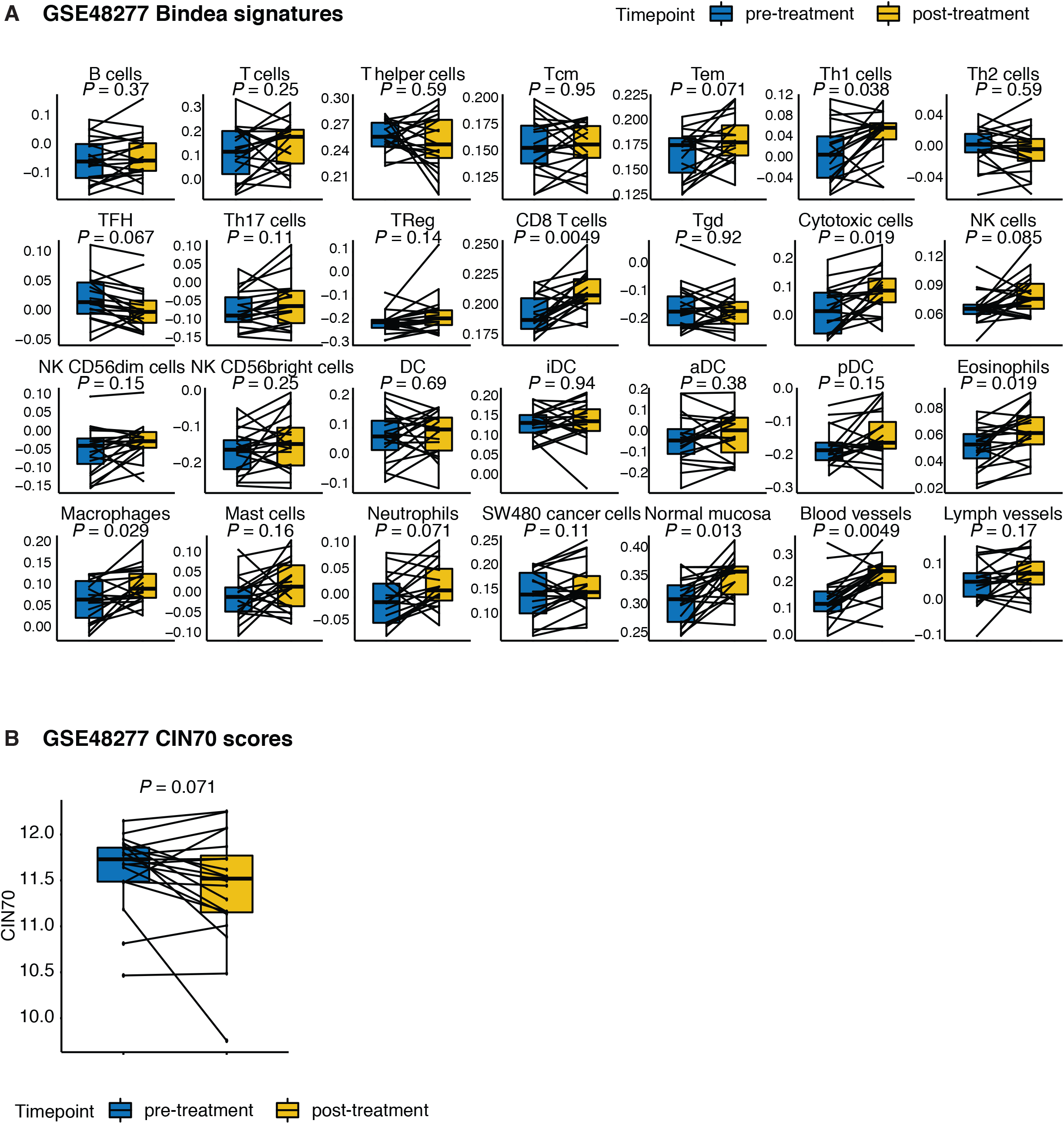
Immune cell signature and CIN70 score changes during bladder cancer treatment. **A)** Paired boxplots showing the Bindea immune deconvolution scores comparing the pre-treatment samples (blue) to the post-treatment samples (yellow) for the bladder cancer dataset GSE48277 (n = 20) in which patients with muscle-invasive bladder cancer received dose-dense neoadjuvant chemotherapy consisting of methotrexate, vinblastine, doxorubicin, and cisplatin prior to radical cystectomy; p-values were calculated using a paired t-test; p-values were corrected for multiple comparisons using the Benjamini-Hochberg method for false discovery rate. **B)** Paired boxplot showing the change in CIN70 score comparing the pre-treatment samples (blue) to the post-treatment samples (yellow) for bladder cancer dataset GSE48277 (n = 20); p-value was calculated using a two-tailed paired t-test.

**Supplementary Table S1.** Patient and tumor characteristics of CRCLM cohorts.

| Variable | UCMC | MSKCC | UK/ New-EPOC | P-value |
| --- | --- | --- | --- | --- |
| Age (mean ± SEM) | 59 ± 1.1 | 60 ± 1.1 | 64 ± 0.9 | 0.0018* |
| Male sex | 58% | 66% | 66% | 0.38 |
| Disease-free interval < 12 months | 55% | 53% | 72% | 0.0028 |
| Maximum liver metastasis size > 5.0 cm | 25% | 23% | 26% | 0.87 |
| Lymph node positive | 64% | 58% | 69% | 0.27 |
| More than 1 liver metastasis | 42% | 60% | 78% | < 0.0001 |
| Elevated CEA (>200 ng/ml) | 4% | 8% | 6% | 0.48 |

P-values were determined using Chi-square tests except \* from one-way ANOVA

